# A novel photoactivatable tool for intermediate filament disruption indicates a role for keratin filaments in early embryogenesis

**DOI:** 10.1101/484246

**Authors:** Rucha Sanghvi-Shah, Shalaka Paranjpe, Jiyeon Baek, Radek Dobrowolski, Gregory F. Weber

**Affiliations:** Department of Biological Sciences, Rutgers University, Newark, NJ 07102, USA; Glenn Biggs Institute for Alzheimer’s & Neurodegenerative Diseases, University of Texas Health Sciences Center, San Antonio, TX, USA; Department of Cell Systems and Anatomy, University of Texas Health Science Center, San Antonio, TX, USA

## Abstract

The significance of cytoplasmic intermediate filament proteins has previously been examined largely through various genetic approaches, including knockdown, knockout and transgenic overexpression. Few studies to date have attempted to examine the role of specifically the filamentous intermediate filament network in orchestrating various cell functions. To directly assess the role of the filamentous keratin intermediate filament network in regulation of cellular behavior, we created a <u>P</u>hoto<u>A</u>ctivatable <u>d</u>isruptor of keratin Intermediate <u>F</u>ilaments (PA-dIF). This genetically encoded construct consists of a peptide derived from the 2B2 region of Keratin 8 fused to the photosensitive LOV2 domain from *Avena sativa* phototropin-1. Upon 458 nm photoirradiation, PA-dIF disrupts keratin intermediate filaments in multiple species and cell types. Marked remodeling of the keratin intermediate filament network accompanies collective cellular morphogenetic movements that occur during gastrulation and neurulation in the *Xenopus laevis* frog embryo. Light-based activation of PA-dIF was able to disrupt keratin intermediate filaments in *Xenopus* cells and lead to tissue-specific disruption of morphogenetic processes. Altogether our data show a fundamental requirement for keratin intermediate filaments in orchestrating morphogenetic movements during early embryonic development that have yet to be revealed in other model systems. Moreover, our data validate the utility of a new genetically encoded photoactivatable tool for the disruption and examination of intermediate filaments.

## Introduction

Intermediate filament proteins polymerize to form cytoskeletal networks present in almost all metazoan cells (Herrmann and Strelkov, 2011). The proteins that constitute this family are expressed in a cell and tissue specific manner. Intermediate filament proteins can broadly be partitioned into two groups based on where they assemble into polymerized filaments. Lamins, the primordial ancestral intermediate filament proteins (Dodemont and Riemer, 1990; Döring and Stick, 1990; Peter and Stick, 2015), assemble into a filamentous network in the nucleus. All other intermediate filament proteins assemble as filaments in the cytoplasm. Cytoplasmic intermediate filaments form five subfamilies that collectively include keratins, vimentin, desmin, neurofilaments, among many others. In addition to their conserved amino acid sequences, motifs and structure, a key defining feature of intermediate filaments is that they convey mechanical strength to cells and cellular structures. Through linkage to cell adhesions, association with the nuclear membrane, and interactions with other cytoskeletal networks, intermediate filaments have important roles in regulating cell shape, nuclear morphology, and consequently cellular function (Sanghvi-Shah and Weber, 2017). The diversity of proteins, and the cytoplasmic filaments that they make, creates small but significant differences in assembly mechanics and the micromechanical properties of filaments (Block *et al.*, 2015).

Keratin filaments polymerize as obligatory heterodimers, requiring both Type I acidic keratin and Type II basic keratin in a 1:1 stoichiometry (Steinert *et al.*, 1976; Hatzfeld and Franke, 1985). These heterodimers associate in anti-parallel to form a tetramer. Tetramers associate laterally with other tetramers to constitute the simplest unit of intermediate filament assembly, the unit length filament (ULF). ULFs associate end-to-end to create a 10-12 nm diameter intermediate filament (Quinlan *et al.*, 1984; Parry *et al.*, 1985; Coulombe and Fuchs, 1990; Steinert, 1990; Martin *et al.*, 2015). This model of the intermediate filament assembly process has largely been derived from *in vitro* work (Herrmann and Aebi, 2016) and comparatively few studies conducted in the cellular context (Helfand *et al.*, 2011; Murray *et al.*, 2014).

Non-epidermal epithelia prominently express Type II Keratin 8 (Krt8/K8) and Type I Keratins 18 and 19 (Krt18/K18 and Krt19/K19) to form K8-K18 and K8-K19 dimers (Franke *et al.*, 1981). Keratin intermediate filaments may then be assembled from a combination of these dimers. Interestingly, Krt8, Krt18, and Krt19, but not vimentin, are also the first cytoplasmic intermediate filaments present and zygotically expressed in the developing embryo (Franz *et al.*, 1983; Suzuki *et al.*, 2017) yet functional roles for these keratin networks remain poorly defined.

Likewise, despite the ubiquity of intermediate filaments, their cellular functions remain poorly understood by comparison to actin microfilaments and microtubules. A variety of tools are readily available for altering actin and microtubule polymerization, which has allowed for major breakthroughs in their fields of research. In this study, we sought to develop a biomolecular tool that could be used to selectively disrupt intermediate filaments.

All intermediate filaments share a common tripartite structure that includes an α-helical rod domain flanked by head and tail regions at both ends. It is through the α-helical rod domain that dimerization occurs to form a coiled-coil. Of particular importance to intermediate filament dimerization is a highly conserved region at the C-terminal end of the coiled-coil domain, termed 2B2 (Hatzfeld and Weber, 1992). Mimetic 2B2 peptide segments incubated with intermediate filament proteins during *in vitro* assembly assays both inhibit the *de novo* assembly of filaments and can induce disassembly of existing filaments. These characteristics seem to be applicable to both vimentin and keratin alike (Kouklis *et al.*, 1992; Steinert *et al.*, 1993b, 1993a; Strelkov *et al.*, 2002). Evidence also indicates that 2B2 peptides can disrupt intermediate filaments in living cells (Helfand *et al.*, 2011).

Advances in optogenetics have enabled the production of genetically encoded proteins that can be activated or otherwise altered by exposure to specific wavelengths of light. The <u>L</u>ight-<u>O</u>xygen-<u>V</u>oltage sensing (LOV)-Jα domain from *Avena sativa* phototropin-1 is one such module that has been used to create various photoswitchable proteins and light-sensitive biomolecular traps (Wu *et al.*, 2009; Lungu *et al.*, 2012; Zimmerman *et al.*, 2016). Here, we have engineered a photoactivatable disruptor of intermediate filaments by fusing the 2B2 domain of Keratin 8 to the LOV-Jα photosensitive module. Using this novel biomolecular tool, we demonstrate the ability to acutely disrupt intermediate filaments with subcellular spatial resolution, and identify important functions for keratin filaments during early embryonic development.

## Materials and Methods

### Plasmid DNA construct design and production

*Xenopus* keratin 8 plasmid C2-eGFP-Krt8 construct was a kind gift from Dr. Victoria Allan and was subcloned into pCS2^+^ previously (Weber *et al.*, 2012). Keratin 19 plasmid pCS2^+^-eGFP-Krt19 was made by Richard Mariani (Mariani *et al.*, 2018). <u>P</u>hoto<u>A</u>ctivatable <u>d</u>isruptor of Intermediate <u>F</u>ilaments (PA-dIF) and <u>C</u>onstitutively <u>A</u>ctive <u>d</u>isruptor of Intermediate <u>F</u>ilaments (CA-dIF) were generated by cloning commercially synthesized Flag-tagged LOV-Jα-2B2 (Flag-PA-dIF) and LOV-Jα(I379E)-2B2 (Flag-CA-dIF) (Genewiz, NJ) into pCS2^+^ vector using EcoRI and XhoI restriction sites. Flag-tag was swapped for mCherry at the N-terminus using NheI and BglII sites in pCS2^+^ Flag-PA/CA-dIF and C1-mCherry.

Plasmid DNA was prepared according to the manufacturer’s protocol using NucleoBond Xtra Midi or Mini kit (Macherey-Nagel). Analytical and preparative restrictions were analyzed by agarose gel electrophoresis (0.8% to 1%) prepared from agarose and 1x TAE running buffer (Tris/Acetate/EDTA: 40 mM Tris-Acetate, pH 8.5, 2 mM EDTA). For visualization of the DNA, either Gel Red (Biotium, #41003) or ethidium bromide (Amersco, #2810) were used and documented using Spectroline UV and UVP PhotoDoc-IT systems.

Purification of DNA fragments from agarose gels or restriction digestions were performed with the Zymoclean Gel DNA Recovery kit (ZymoResearch, #D4002) or Nucleospin Gel and PCR clean up (Macherey-Nagel, #74060950) according to the manufacturer’s instructions.

Concentration of restriction-digested and gel purified vector and insert DNA fragments was determined by A260 on a spectrophotometer. 1:3 to 1:10 molar ratio of vector:insert was used in ligation reactions. Standard ligation reactions were carried out using T4 DNA Ligase (New England Biolabs, #MO202L) according to the manufacturer’s instructions. The ligation was incubated for 45 minutes at room temperature or overnight at 15°C.

For chemical transformation of the ligation reaction, 5-alpha (New England Biolabs, #C2988J) or Express Iq competent *E. coli* cells (New England Biolabs, #C3037I) were used. The ligation reaction volume was added to 50 μl of the competent cells in a pre-chilled 14 ml round bottom tube on ice and incubated for 30 minutes. Cells were heat shocked at 42°C for exactly 2 min, followed by a further incubation on ice for 2 minutes. SOC medium (1 ml) was added and the transformed cells were incubated for 60 min at 37°C in a rotating shaker (∼250 rpm) for recovery. Subsequently, different quantities of transformed cells were spread onto warm selection plates (LB agar plates supplemented with the appropriate antibiotic; ampicillin or kanamycin) and colonies were allowed to grow overnight at 37°C.

### Software analyses of protein structure

Protein sequences were analyzed for hydrophobic moment and amphipathicity using Heliquest (http://heliquest.ipmc.cnrs.fr/) (Gautier *et al.*, 2008). Predicted secondary structure and 3D models of protein constructs were generated using PHYRE^2^ (http://www.sbg.bio.ic.ac.uk/phyre2/html/page.cgi?id=index) (Kelley *et al.*, 2015).

### *In vitro* transcription

To synthesize RNA for microinjection, 6 µg of pCS2^+^-PA-dIF, pCS2^+^-CA-dIF, or pCS2^+^-EGFP-Krt8 DNA was linearized with NotI restriction enzyme (New England Biolabs) at 37°C overnight. pCS2^+^-EGFP-Krt19 DNA was linearized with BssHII (New England Biolabs) at 50°C overnight. Digested DNA was extracted with phenol:chloroform:isoamyl alcohol (25:24:1) (Invitrogen) and precipitated with 3 M sodium acetate solution and 100% ethanol and incubated overnight at - 20°C. DNA was pelleted by centrifugation at 14000 g for 10 min at 4°C and washed once using 100 µl 70% ethanol and centrifuged for 5 min, 14000 g at 4°C. Ethanol was removed and the pellet was dried at 37°C. DNA was then solubilized in 16.5 µl nuclease free water. *In vitro* transcription was performed in a reaction mixture containing 5 µl 10x transcription reaction buffer, 5 μl 100 mM rATP, rCTP, rUTP ribonucleotide mix (Promega, #P1420) 5 μl 1 mM rGTP, 5 μl 10 mM m7G(5’)G RNA Cap (New England Biolabs, #S1404S), 1 μl RNase inhibitor (Promega, #P1420), and SP6 RNA polymerase (Promega, #P1420) and incubated at 37°C for 30 minutes. 1.25 µl of 10 mM rGTP was added and incubated for additional 1 hour at 37°C. To avoid degradation of the RNA, 0.5 μl of the RNase inhibitor and 2.5 μl RQ DNase1 (Promega, #P1420) were added followed by incubation for 15 minutes at 37°C. Free nucleotides and digested DNA were removed using Illustra ProbeQuant G-50 Microcolumns. Synthesized RNAs were purified by phenol:chloroform extraction and ethanol precipitation as above and stored at - 80°C.

### *Xenopus laevis in vitro* fertilization and embryo culture

Adult *Xenopus laevis* were obtained from Nasco (Fort Atkinson, WI) and housed at Rutgers University-Newark and used in accordance with institutional guidelines and the approval of the local Institutional Animal Care and Use Committee. Female frogs were injected subcutaneously with a priming dose of human chorionic gonadotropin (hCG) 50 U (2 U/μl) (MP Biomedicals, #198591) in the dorsal lymph sac 7-10 days preceding induction of ovulation. Females were induced to ovulate by injection of 500 U human chorionic gonadotropin and incubated at 15°C until ovulation commenced, typically 12 hours later. Females were manually ‘squeezed’ to yield eggs.

To achieve synchronous fertilization, ∼1 mm thick fragment of the testis was macerated and further homogenized in 1 ml 1x Modified Barth’s Saline (1x MBS: 88 mM NaCl, 1.0 mM KCl, 2.4 mM NaHCO_3_, 15.0 mM HEPES (pH 7.6), 0.3 mM Ca(NO_3_)_2_-4H_2_O, 0.41 mM CaCl_2_-2H_2_O, 0.82 mM MgSO_4_). Eggs were thoroughly mixed with the sperm suspension and allowed to sit for 3 minutes before flooding the Petri dish with deionized water. After 20 mins fertilized eggs were dejellied by gently swirling (3-5 minutes) in a 2% cysteine hydrochloride (Amresco, #200-157-7), pH 7.8 solution in 0.1x MBS. Dejellied zygotes were thoroughly rinsed with deionized water followed by 0.1x MBS. Zygotes were then cultured in 0.1x MBS at 15°C until they were ready to be used for experiments.

### Embryo microinjections

To target ectodermal (animal cap) tissue, *Xenopus* zygotes were microinjected at one-cell stage in the animal hemisphere. 5 ηl of appropriate DNA (Krt8/Krt19, 15 pg/ηl; CA-dIF/PA-dIF, 50 pg/ηl) or RNA (Krt8/Krt19, 100 pg/ηl; CA-dIF/PA-dIF, 20-50 pg/ηl) diluted in deionized water was pressure injected using a Narishige IM-300 microinjection apparatus. Microinjection needles were fabricated from borosilicate glass capillaries (Drummond, #1000 0010) using a P-30 vertical micropipette needle puller (Sutter Instrument, CA) at settings 760 units for heat and 400 units for pull. Needles were calibrated to the desired injection pressure (20-30 PSI) and time (100-200 ms) settings to dispense 5 ηl of injection volume. During injections and 20 minutes post injection, the embryos were kept in 3% Ficoll (Amresco, #E965-504). Embryos were rinsed thoroughly and transferred to 0.1x MBS and allowed to develop to the gastrulation stage by maintaining in a 15°C incubator. To avoid unintended photoactivation, embryos were maintained in the dark and all experimental steps were performed under red light only.

### Preparation of ectodermal tissue (**animal caps**) explants

Basic animal cap explants were prepared from *Xenopus* embryos at gastrulae stage 11-11.5 in a 100 mm dish containing 0.1x MBS. First, the embryos were devitellinized from the vegetal side using forceps to avoid damaging the animal cap. Next, the animal caps were excised using the eyebrow knife. The blastocoel side of the animal hemisphere was exposed from the vegetal side and then the region of the animal cap with fluorescence was identified and cut. To obtain the animal cap region expressing the injected proteins, the sides of the explants were shaved to appropriate size. The explanted animal caps were then transferred to a 35 mm dish coated with fibronectin (1.4 µg/cm^2^) containing 0.5x MBS using a Pasteur pipette. Explants were positioned with the deep layer facing the bottom of the chamber. After arranging the explant on the substrate, a small fragment of the cover slip supported by silicone grease at the four corners was used to compress the explant gently with forceps. Ectodermal explants were then allowed to attach onto the substrate for about 30 minutes before image acquisition.

### Developmental phenotypes

Synchronously fertilized embryos were harvested for analysis at the desired developmental stage according to the external morphology as described by (Nieuwkoop and Faber, 1994). Embryos were kept at a density of maximum 50 embryos per 35 mm dish and 0.1x MBS was replaced daily.

### Cell culture and transfection

Madin-Darby canine kidney epithelial cells and HEK293T cells were obtained from ATCC (CCL-34 and CRL-3216, respectively). Cells were maintained in growth medium consisting of Dulbecco’s modified Eagle’s medium (DMEM) (Gibco, #31053028) containing 10% Fetal bovine serum (FBS) (Gibco, #10438026), 1% Penicillin-Streptomycin (Gibco, #P0781), 1% Glutamax (Gibco, #35050-061), 26.18 mM NaHCO^3^ and filtered sterilize using 0.2 µm vacuum filter. Cells were grown at 37°C with 5% CO^2^ in a humidified tissue culture incubator. Cells were not used beyond passage 20 and were cultured from frozen stock.

HEK293T and MDCK cells expressing mCherry CA-dIF/PA-dIF were generated by transfection of the pCS2^+^ mCherry-PA/CA-dIF construct using BioT plasmid transfection reagent (Bioland Scientific, B01-00) or Lipofectamine 3000 (Invitrogen, Life Technologies), respectively, according to manufacturer’s guidelines. Briefly, cells were seeded onto 35 mm dishes to achieve 70-80% confluency at the time of transfection. Next, 10 µl of Opti-MEM reagent was incubated with 0.3 µl of Lipofectamine, 0.2 µl of P3000 and 0.1-0.2 µg of DNA for 15 minutes and then the DNA-lipid complexes were added on to the seeded cells. For 35 mm dishes, 150 µl of Opti-MEM reagent was incubated with 10 µl of Lipofectamine, 7.5 µl of P3000 and 1-1.5 µg of DNA for 15 minutes and then the DNA-lipid complexes were added on to the seeded cells. Cells were incubated for 6 hours at 37°C before exchanging for fresh cell culture medium. Cells transfected with PA-dIF constructs were handled in dark or under red light to avoid unintended activation.

### *Xenopus laevis* embryo lysis and biochemistry

Embryos at stage 10.5-11 were sorted for eGFP or mCherry fluorescence to ensure expression of proteins from the injected mRNA. Fluorescence-positive embryos were homogenized on ice with pre-chilled extraction buffer (100 mM NaCl, 50 mM Tris-HCl (pH 7.5), 1% Triton X-100, 1 mM phenylmethylsulfonofluoride (PMSF), 10 mg/ml sodium β-glycerophosphate, 10 mM sodium fluoride, 1 mM sodium orthovanadate, 0.2 mM H_2_O_2_, 3 mM sodium pyrophosphate, with mammalian protease inhibitor cocktail (Sigma, #P2714)). The embryo slurry was incubated on ice for 15 mins followed by centrifugation for 10 minutes at 14,000g at 4°C. To clarify the lysate, the yolk from the upper phase was aspirated. If a sequential aspiration was necessary the supernatant was centrifuged again with the same conditions. If the lysate was not used immediately, samples were frozen at -80°C, aliquoted for SDS/PAGE, and/or immunoprecipitated.

#### Co-immunoprecipitation

Immunoprecipitations were performed with RFP-Trap_MA (Chromotek, Germany, #rtma20) using 100 μl total lysates extracted from uninjected embryos or embryos injected with either eGFP-Krt19 (100 pg/ηl) alone or eGFP-Krt19 and mCherry PA-dIF (100 pg/ηl). Reactions were incubated for 1 hour at 4°C, beads were then magnetically separated and further washed 3 times with lysis buffer. Immunocomplexes were dissociated from beads by 40 μl of 2x Laemmli buffer with 5% β-mercaptoethanol and boiling for 10 minutes. Samples were then separated by SDS-PAGE. Polyacrylamide gels of 12% or 14% concentration were prepared by mixing appropriate volumes of 1.5 M Tris-HCl pH 8.8, 30% Acrylamide-Bis acrylamide (Biorad), 10% SDS, 10% ammonium persulfate (Sigma), 0.02% TEMED (VWR). A 4% stacking gel was made by mixing appropriate volumes of 0.5 M Tris-HCl pH 6.8, 30% acrylamide (Biorad), 10% SDS, 10% ammonium persulfate (Amresco), 0.02% TEMED (VWR). Gels were loaded with protein samples and run at 80-100V. After separation, proteins were stained with Sypro Red dye (Invitrogen, S12000).

#### LC/MS-MS Proteomic Analysis

Prior to staining with Sypro Red dye, SDS-PAGE gels were incubated twice in fixative solution consisting of 50% methanol and 7% glacial acetic acid for 30 minutes per incubation. After decanting the second fixative solution, the gel was placed in a fresh dish and incubated in 60 ml of Sypro Red dye overnight. The staining solution was decanted and the gel was incubated in a wash solution of 10% methanol and 7% glacial acetic acid for 30 minutes. Afterwards, the gel was washed three times in 100 ml of commercial ultrapure water before gel imaging and further preparation for LC/MS-MS. All incubations and washes were performed at room temperature on a flat rotator.

LC/MS-MS was performed by the Center for Advanced Proteomics Research (CAPR) at the Rutgers New Jersey Medical School. Sypro Red-labeled SDS-PAGE gel sections were excised at the facility and in-gel trypsin digestion was performed. The resulting peptides were C18 desalted and analyzed by LC/MS-MS on the Q Exactive instrument. The MS/MS spectra were searched against the NCBI *Xenopus laevis* database using Sequest search engines on the Proteome Discoverer (V2.1) platform. The protein false discovery rate (FDR) is less than 1%. The comparison of identified proteins and their relative quantitation was calculated based on the spectra counting method and compared for consistency across two independent experiments. In addition to the 1% FDR, for inclusion in the final data table shown in Figure 3, proteins had to meet all of the following criteria: molecular weight between 40-75 kDa, 3 or more unique peptides in each single-run analysis, spectral count in the CA-dIF sample of 10 or more, and a ratio of enrichment in the CA-dIF sample of 1.5 or greater.

### Indirect Immunofluorescence

MDCK or HEK293T cells were seeded on 35 mm glass bottom dishes (MatTek). Cells were fixed in ice cold 100% methanol for 10-15 mins on ice and then washed 3 times with PBS. Cells were permeabilized with 0.25% Triton X-100 diluted in PBS and rinsed 3 times with PBS. Cells were blocked for 1hr at 37°C with 10% goat serum, 1% BSA and 0.1% Triton X-100 diluted in PBS, following which samples were incubated with primary antibodies diluted in blocking buffer for 30 mins at 37°C or overnight at 4°C. Samples were then thoroughly washed with PBS and then incubated with secondary antibodies, Hoechst 34580 (1:10,000; Sigma Aldrich, #63493) for 30 mins at 37°C, rinsed again 3 times with PBS.

Pan-cytokeratin (1:100, C2562, Sigma-Aldrich) primary antibody was used for immunofluorescence. Secondary antibodies used were species-specific AlexaFluor-IgG conjugates (1:1000) (Life Technologies). mCherry fluorescence was preserved through fixation and labeling.

### Confocal microscopy and image analysis

Confocal images were acquired with Zeiss Cell Observer SD using Zen software (Zeiss) equipped with a Plan-Apochromat 63X/1.4 oil immersion objective, 1.0 or 1.6 Optovar. Z-stacks with a step size of 0.25-0.30μm were acquired for each field. For optimal images, z-stack maximum intensity projections were processed from z-stacks collected through the entire cell and only linear adjustments were made to their brightness and contrast.

Linescan measurements of fluorescence were conducted using the linescan tool in the profile tab of the Zen 2.3 lite software application. Measurements were taken in a region of cytoplasmic volume for a length of 15 μm. Intensities were normalized to the intensity value at the beginning of the linescan in order to view intensity fluctuation.

For photoactivation studies, embryos were co-injected with mCherry PA-dIF or mCherry CA-dIF (DNA, 50 pg/ηl) and eGFP-Krt8 (DNA, 15 pg/ηl). Ectodermal explants were prepared as described above. Because we observed that high expression of PA-dIF may disrupt keratin networks even when maintained in dark conditions, cells with apparently normal IF networks and general cell morphology and having moderate expression of mCherry PA-dIF were selected for photoactivation. Cells were imaged with a Zeiss Cell Observer SD confocal microscope using a Plan-Apochromat 63X/1.4 oil immersion objective, 1.0 Optovar and a DirectFRAP module. To induce PA-dIF activation, a 458 nm laser was focused into a small circular area (diameter 10 μm) for 30 seconds with 100% laser intensity. This combination of settings yielded an 8 μW blue light exposure. Images were acquired every 15 secs for 1 min before and between irradiation periods, and then for 5-10 minutes following the last irradiation.

### Photoactivation of PA-dIF in embryos

For blue light stimulation of whole embryos, 35 mm dishes containing either control or mCherry PA-dIF (CA-dIF/PA-dIF, 20-50 pg/ηl) injected embryos were placed underneath a home-built (50X40 mm) LED array. Briefly, an LED array was constructed with 56 blue 5 mm LEDs (470 nm, 7065 millicandelas each max output) with a mean 100 μW measured output. The LEDs were powered through a 40 V power supply with a potentiometer for intensity control. Following microinjection of mCherry PA-dIF, the embryos were illuminated with oscillating pulses every sec with a minimum 53 μW and maximum of 154 μW intensity or maintained in the dark, until the embryos reached desired stages. Embryos were fixed in 4% formaldehyde at desired timepoints and analyzed for morphology.

## Results

### Rational design of a photoactivatable disruptor of keratin intermediate filaments

The highly conserved C-terminal portion of the α-helical rod domain, 2B2, is known to be critical for the dimerization of intermediate filaments (Fig. 1A) (Strelkov *et al.*, 2002). Previous work has shown that as few as 20 amino acids from this region is sufficient to disrupt keratin assembly *in vitro* (Hatzfeld and Weber, 1992; Steinert *et al.*, 1993b, 1993a). Dimerization of these mimetic peptides with their endogenous keratin partner prevents further assembly into mature filaments, resulting in a combination of intermediate filament precursors and more general filament disruption and aberrant aggregation. Similarly, peptides resembling the 2B2 region of intermediate filaments efficiently disrupt intermediate filaments *in vivo* (Helfand *et al.*, 2011).

**Figure 1:**
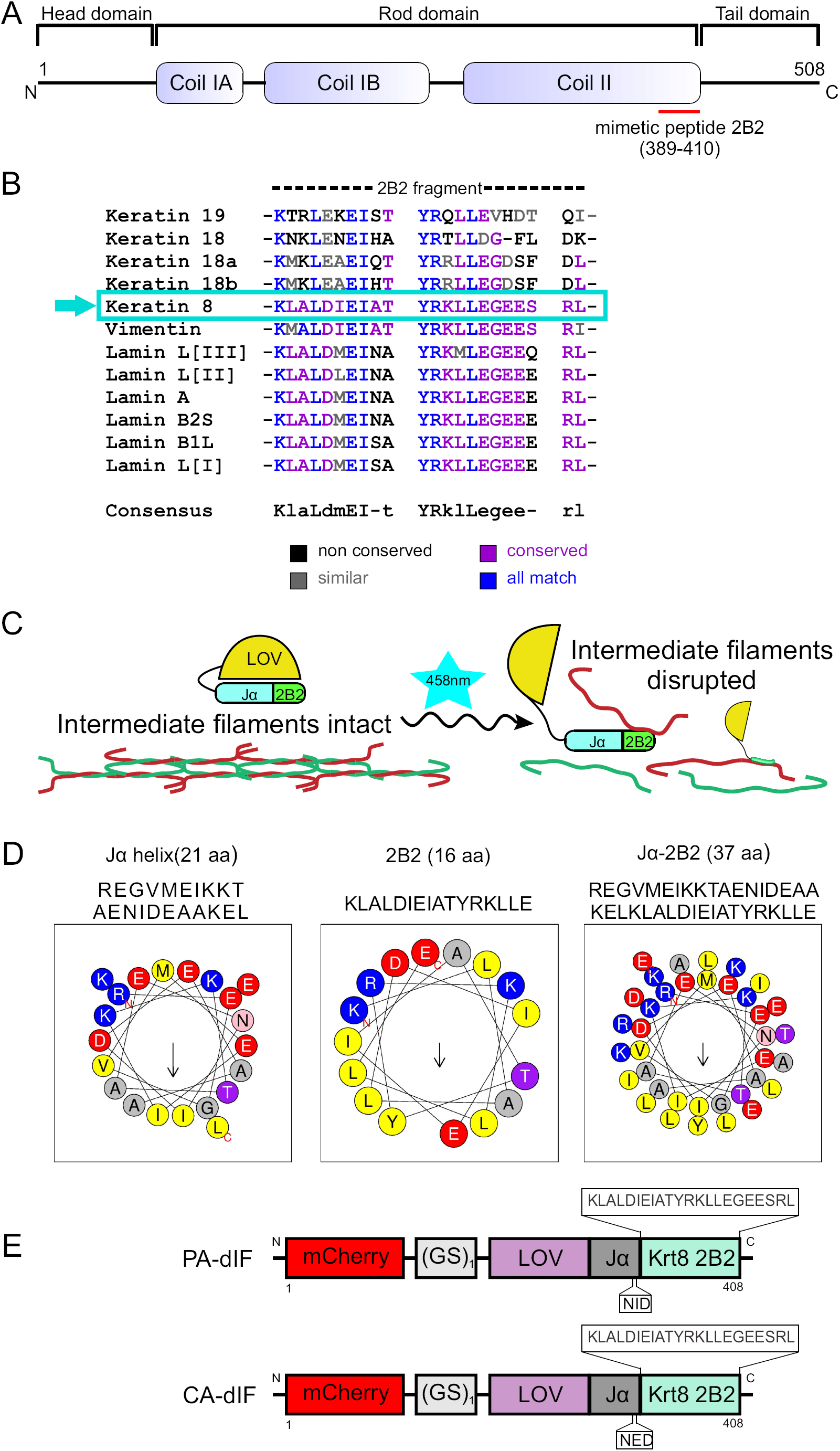
Design of photoactivatable disruptor of keratin intermediate filaments. **(A)** Schematic representation of primary structure of keratin IF proteins. The central rod domain, comprised of α-helical coils IA, IB and coil II, are flanked by variable length head and tail domains at the N- and C-termini, respectively. The red bar denotes the segment of keratin used in the constructed fusion proteins. (**B)** Sequence alignment of 2B2 segments of *Xenopus laevis* IF proteins including keratin 8, keratin 18/19, vimentin and nuclear lamins A, B and L. High homology exists across intermediate filaments. Keratin 8 2B2 has particularly high homology with vimentin and lamins. (**C)** Strategy for caging 2B2 mimetic peptide and mechanism for keratin network disruption. Upon irradiation with 458nm light the LOV-Jα is predicted to uncouple, making the 2B2 peptide available to bind to its dimerization partners keratin 18/19 and thus disrupt keratin filaments. (**D)** Conserved amphipathic pattern in Jα-helix facilitates interactions between 2B2 and Jα-helix. Helical wheel projection of Jα-helix, 2B2 and 2B2 appended to Jα-helix shown here were generated using Heliquest software molecular modeling. (**E)** Schematics of PA-dIF and CA-dIF modules comprise a *Xenopus laevis* Krt8 2B2 (22aa) mimetic peptide which is directly fused to the LOV-Jα *Avena sativa* photosensitive domain by a GS linker. CA-dIF module has an I379E point mutation in the Jα domain.

We sought to create a fluorescently-traceable photoactivatable fusion protein that contained 2B2 and which could be acutely ‘activated’ to disrupt intermediate filaments with subcellular spatial resolution. Based on data from Hatzfeld and Weber (Hatzfeld and Weber, 1992), we used a near-minimum number of amino acids from the C-terminal part of 2B2 derived from *Xenopus* Krt8 as the template for our inhibitory peptide. This amino acid sequence is highly conserved across intermediate filament proteins (Fig. 1B) and species (Fig. S1). 2B2 was fused to the Jα helical region of the LOV-Jα domain from *A. sativa* phototropin-1 (Fig. 1C, Fig. S2). Several strategies noted by other laboratories were used in our design in order to maximize caging while allowing 2B2 to be available for binding upon photoirradiation. Tighter caging was suggested to be enhanced by minimizing peptide sequence that extends beyond the LOV-Jα interface. Both Jα and 2B2 are amphipathic α-helices (Fig. 1D). The hydrophobic side of the Jα helix folds against the adjacent globular LOV domain (Harper *et al.*, 2003; Möglich *et al.*, 2009). In the case of 2B2, coiled-coil formation of the intermediate filament dimer is facilitated by this amphipathic property. In order to promote α-helix formation and folding of the fusion helix against the globular LOV region, we performed a series of *in silico* analyses. Lungu et al. suggested that embedding the peptide within the Jα helix yields improved caging potential but with the caveat that accessibility of the peptide may be limited, even in the lit state (Lungu *et al.*, 2012). In an effort to balance this cost-benefit, we fused the 22 amino acid 2B2 sequence directly to the end of Jα. Fusion of 2B2 to Jα is predicted to create a nearly contiguous α-helical domain (Fig. S3A,B). Analysis of the α-helix formed by Jα and 2B2 showed that the amphipathic nature of the helix was predicted to be preserved in a fusion construct (Fig. 1D). A continuous α-helix with a similar hydrophobic moment to either Jα or 2B2 alone is expected. Predictive 3D modeling of sequences encoding LOV-Jα, LOV-Jα-2B2, and mCherry-LOV-Jα-2B2 were performed using Phyre2 (Kelley *et al.*, 2015). Modeling of LOV-Jα closely resembles other predicted and actual X-ray crystallography- and solution NMR spectroscopy-based structures of the domain (Fig. S4A) (Harper *et al.*, 2003; Wu *et al.*, 2009; Lungu *et al.*, 2012). The addition of the intermediate filament 2B2 domain extended the Jα helix (Fig. S4B). Interestingly the addition of a mCherry fluorophore to the fusion protein generated a loop and turn in the Jα-2B2 helix (Fig. S4C). In either case, the 2B2 domain is predicted to be helical in structure and associate with the globular LOV domain. An additional construct containing a mutation within the Jα domain of I379E yielded a ‘lit-state’ variant (Fig. 1E) that cannot fold properly due to steric hindrance of the LOV-Jα interaction (Fig. S4D) (Harper *et al.*, 2004; Wu *et al.*, 2009; Zimmerman *et al.*, 2016). These strategies yielded two constructs: 1) a <u>P</u>hoto<u>A</u>ctivatable <u>d</u>isruptor of Intermediate <u>F</u>ilaments (PA-dIF) and 2) a <u>C</u>onstitutively <u>A</u>ctive <u>d</u>isruptor of Intermediate <u>F</u>ilaments (CA-dIF).

### CA-dIF perturbs intermediate filament network and nuclear morphology

We first sought to test these constructs for their caging ability and whether the 2B2 sequence would still remain functional after having been fused with the Jα. Although the sequence of 2B2 used originates from frog Krt8, the high homology suggested that this sequence should disrupt keratin intermediate filaments across species and cell types (Fig. S1). We transfected HEK293T and MDCK cells with either CA-dIF or PA-dIF to test the ability of the constructs to disrupt IFs and the ability of the LOV-Jα domain to effectively mask the 2B2 peptide. Transfected cells were maintained in typical incubator conditions in the dark where PA-dIF would not be activated. HEK293T cells transfected with PA-dIF and maintained in the dark exhibited normal spindly keratin networks typical for this cell line (compare untransfected mCherry-cells to transfected mCherry+ cells, Fig. 2A). Nuclei in these cells were similarly robust and oval in shape. In contrast, HEK293T cells expressing CA-dIF showed a punctate perinuclear aggregation of the keratin network (Fig. 2B). Nuclei in CA-dIF cells were often lobulated and dysmorphic.

**Figure 2:**
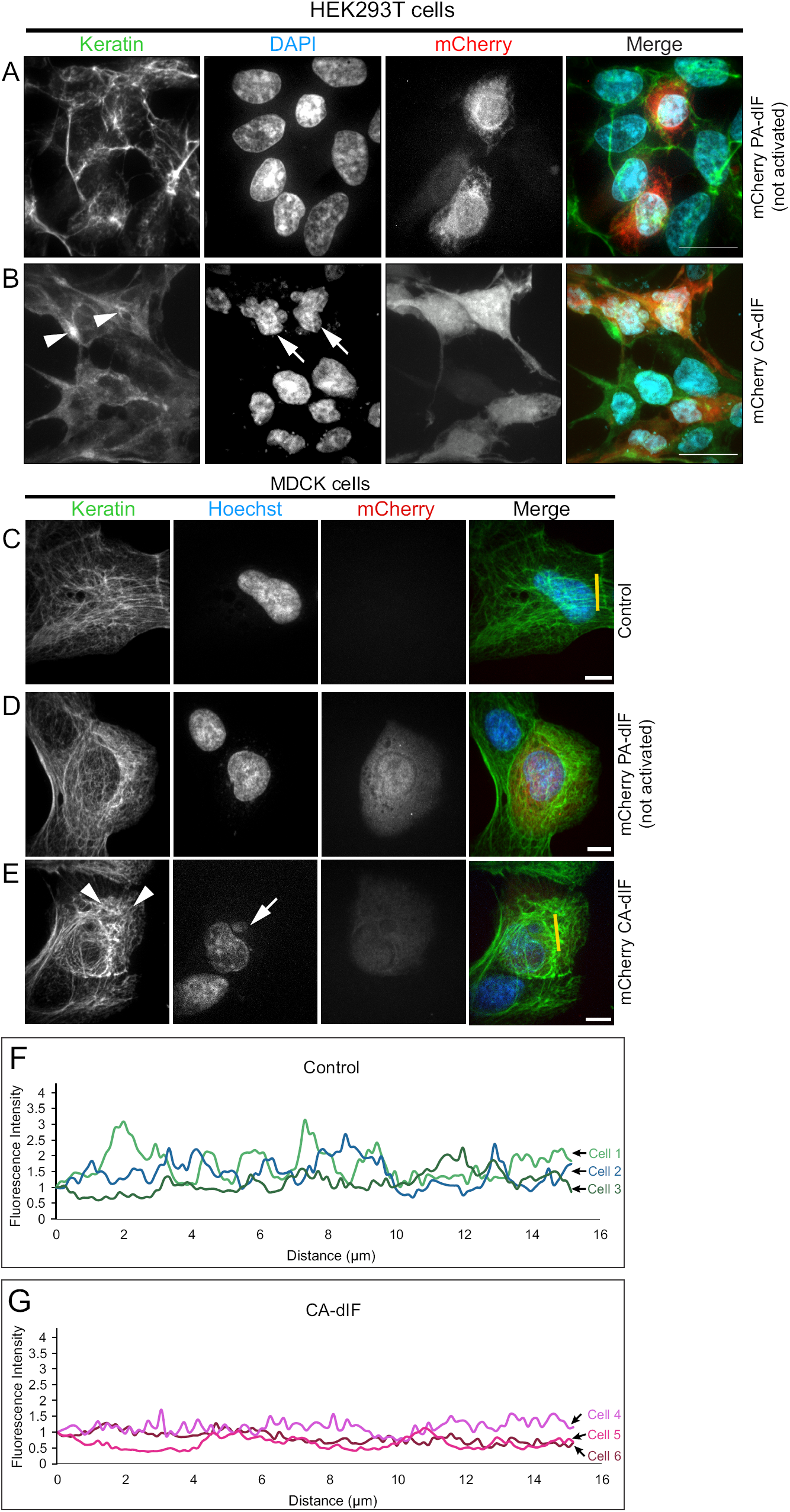
Constitutively active disrupter of intermediate filaments (CA-dIF) disrupt keratin filaments in cultured HEK293T cells and MDCK cells. The keratin network (green) was examined in HEK293T cells transfected with either mCherry PA-dIF (**A)** or mCherry CA-dIF (**B)**, 24 hrs post-transfection. Cells were immunostained with pan-keratin antibody and counterstained with DAPI. Bars, 20 μm. MDCK cells were either mock transfected (control) (**C)** transfected with mCherry PA-dIF (**D)** or mCherry CA-dIF (**E)**, 12 hrs post transfection. Confocal images of cells immunostained with pan-keratin antibody and counterstained with Hoechst dye display keratin network (green) organization and expression of PA-dIF and CA-dIF constructs (red) and nucleus (blue). Bars, 10 μm. White arrowheads show disruption of keratin filaments and white arrows show aberrant nuclear morphology. Yellow bar line in merged image indicates example region of line scans shown in graphs in (**F)** and (**G)**. Distinct filaments are seen in control line scans as evidenced by periodic intensity fluctuations (**F)**, whereas keratin filaments in mCherry CA-dIF transfected cells are more aggregated and/or less distinct as shown by less varied intensity fluctuations (**G)**.

MDCK cells originated from canine kidney epithelia and are enriched with a diverse number of keratin intermediate filament proteins. Normal untransfected MDCK cells have a well-spread keratin network distribution and oval nuclei (Fig. 2C). No observable differences were noted between untransfected MDCK cells and mCherry PA-dIF cells (Fig. 2D). MDCK cells transfected with CA-dIF showed perinuclear clustering of the keratin network, although not to the same degree as HEK293T cells (Fig. 2E). Filaments were shorter and less distinct, which produced less intensity fluctuation by line-scan analyses of keratin labelling in the cytoplasm (Fig. 2F,G). Like HEK293T cells, MDCK cells also showed some nuclear morphology changes, but again, these were not as severe. Altogether, these data show the capacity of CA-dIF to disrupt keratin intermediate filaments in multiple cell and species types. In addition, these findings provide evidence that suggests PA-dIF is effectively caged when maintained in the dark.

### Photoactivation of PA-dIF disrupts keratin intermediate filaments

While peptides of truncated 2B2 region of intermediate filaments have been demonstrated to disrupt intermediate filaments *in vitro*, these peptides have not been widely used to perturb intermediate filaments in cells and, to our knowledge, no studies to date have used this peptide *in vivo* in the whole organism. Previously, peptide mimetics of the 2B2 domain have been used in *in vitro* assays of intermediate filament assembly and in a few studies microinjected into living cells (Hatzfeld and Weber, 1992; Steinert *et al.*, 1993a; Strelkov *et al.*, 2002; Helfand *et al.*, 2011). In principle, by caging this peptide with the photoactivatable LOV-Jα module, our developed tool could be light-activated in living cells to disrupt intermediate filaments at the subcellular spatial scale.

Early *Xenopus laevis* embryos provide a useful model organism in which to investigate the developmental and cell biological roles of intermediate filaments. During early embryonic development, expression of intermediate filament proteins is limited to Krt8, Krt18, and Krt19 into the early morphogenetic events of gastrulation with vimentin emerging in the ectodermal layers toward late gastrulation. The Krt8/18/19 keratin complement is present in all tissues of the embryo from egg through gastrula. Fluorescently-tagged Krt8 or Krt19 readily incorporates into intermediate filament networks of the *Xenopus* embryo and can be used to track the keratin network in living tissue (Weber *et al.*, 2012; Mariani *et al.*, 2018). To investigate the ability of dIF constructs to disrupt keratin networks in an *in vivo* model system, we co-injected mRNA encoding PA-dIF or CA-dIF with either eGFP-Krt8 or eGFP-Krt19 into the animal cap of *Xenopus* embryos. Ectodermal animal cap explants were prepared at gastrulation and keratin intermediate filaments were imaged in live cells of the deep cell layers. Keratin intermediate filament networks were distributed throughout the cell body of ectodermal cells (Fig. 3A), similar to that previously described in unpolarized mesendoderm cells (Weber *et al.*, 2012). Likewise, coalescence of keratin filaments was seen occasionally near cell-cell contacts.

**Figure 3:**
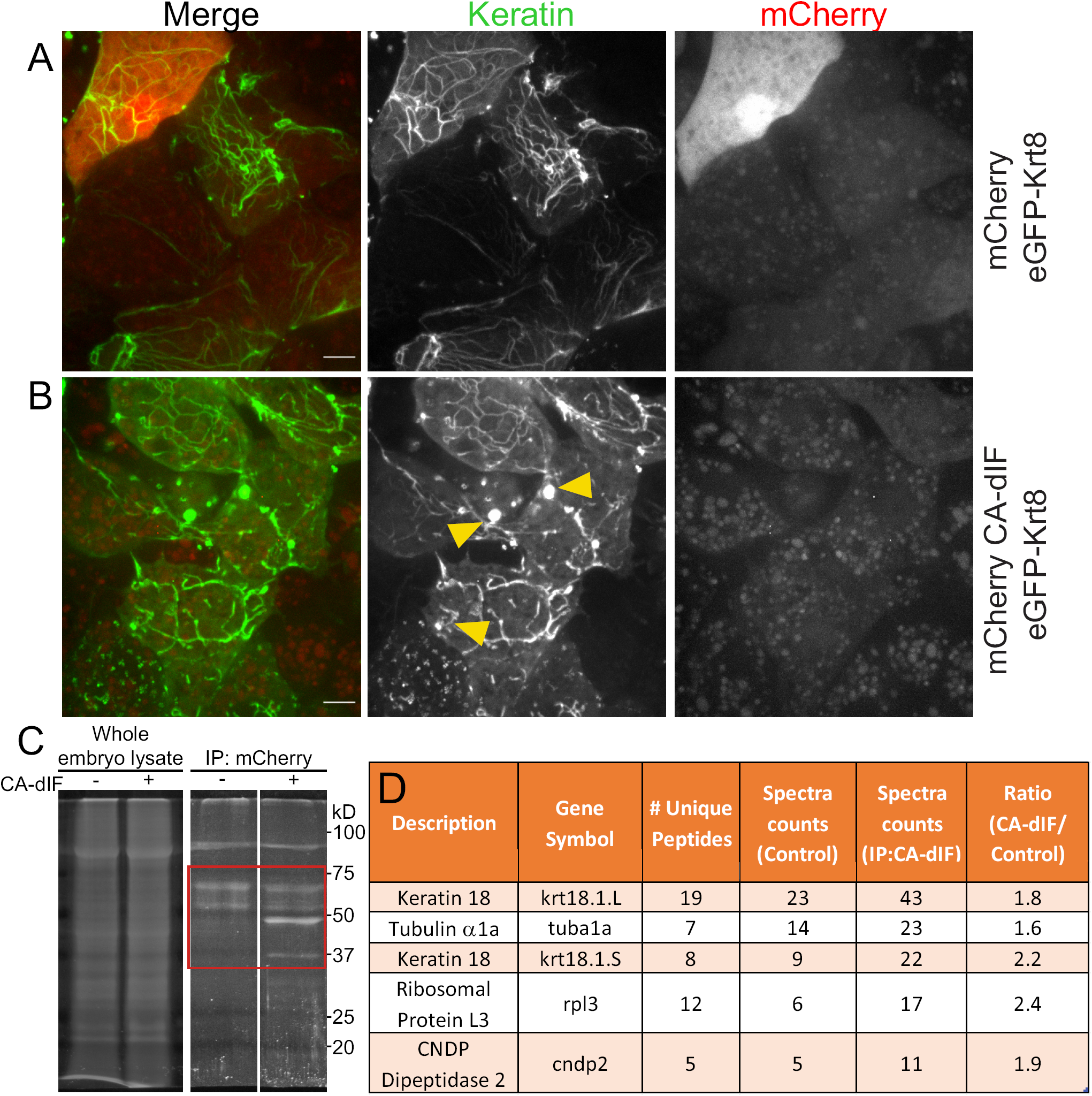
Expression of CA-dIF binds to and disrupts keratin filaments in embryonic *Xenopus* cells. Representative confocal images of *Xenopus* ectodermal explants (stage 11) expressing eGFP-Krt8 and either mCherry (**A),** or mCherry-CA-dIF (**B)**. Yellow arrowheads indicate the disruption of the keratin filaments in cells expressing mCherry-CA-dIF even at low levels. Bars, 10 μm. *Xenopus* embryos were microinjected with mCherry-CA-dIF (500 pg) mRNA at one cell stage in the animal hemisphere. (**C)** Lysates from embryos in gastrula stage were subjected to immunoprecipitation to isolate mCherry-CA-dIF complexes and separated by SDS-PAGE. Protein bands within the region outlined by the red box were excised, in-gel trypsinized and analyzed by LC-MS/MS. (**D)** Table summary of abundant proteins detected in LC-MS/MS samples.

We first co-expressed mCherry-CA-dIF and eGFP-Krt8 to examine whether the filamentous keratin network was disrupted when the open-state fusion construct was expressed (Fig. 3B). Keratin filaments formed punctate aggregates throughout mCherry positive cells, even at low expression levels. To confirm the molecular target of our dIF constructs when expressed *in vivo*, we immunoprecipitated CA-dIF from injected *Xenopus* gastrula and looked for enrichment of proteins in mCherry-CA-dIF samples (Fig. 3C). Protein bands were enriched in the molecular weight range of 37-75kD. These bands were isolated and liquid chromatography-tandem mass spectroscopy analyses were performed to identify endogenous proteins that were binding to CA-dIF. Keratin 18 isoforms originating from both *X. laevis* alleles predominated in terms of both unique peptides identified, spectral counts, and were identified as the top target proteins of our construct with few other proteins enriched (Fig. 3D).

To test the ability of PA-dIF to disrupt intermediate filaments, cells expressing PA-dIF were photoirradiated with 458 nm light for a period of 30 seconds while performing time-lapse imaging. No change to the intermediate filament network was observed in cells expressing eGFP-Krt8 and mCherry (Fig. 4A). In cells expressing PA-dIF, exposure to blue light induced the rapid collapse of keratin filaments specifically in the immediate vicinity of illumination (Fig. 4B-D). While total collapse of the keratin filaments was isolated to the region of photoirradiation, this localized perturbation had widespread consequences for the intermediate filament network beyond the local effect, causing filaments nearby to appear more tortuous rather than taut. However, only in the area where illumination occurred did filaments completely collapse and aggregate. To further explore the capability of PA-dIF to disrupt the intermediate filament network in an acute manner, we performed repeated illuminations of a single cell where we systematically targeted filaments that remained until the entire intermediate filament network was collapsed. Repeated photoirradiation was able to acutely disrupt the entire intermediate filament network within a cell in a period of less than 10 minutes (Fig. 4D).

**Figure 4:**
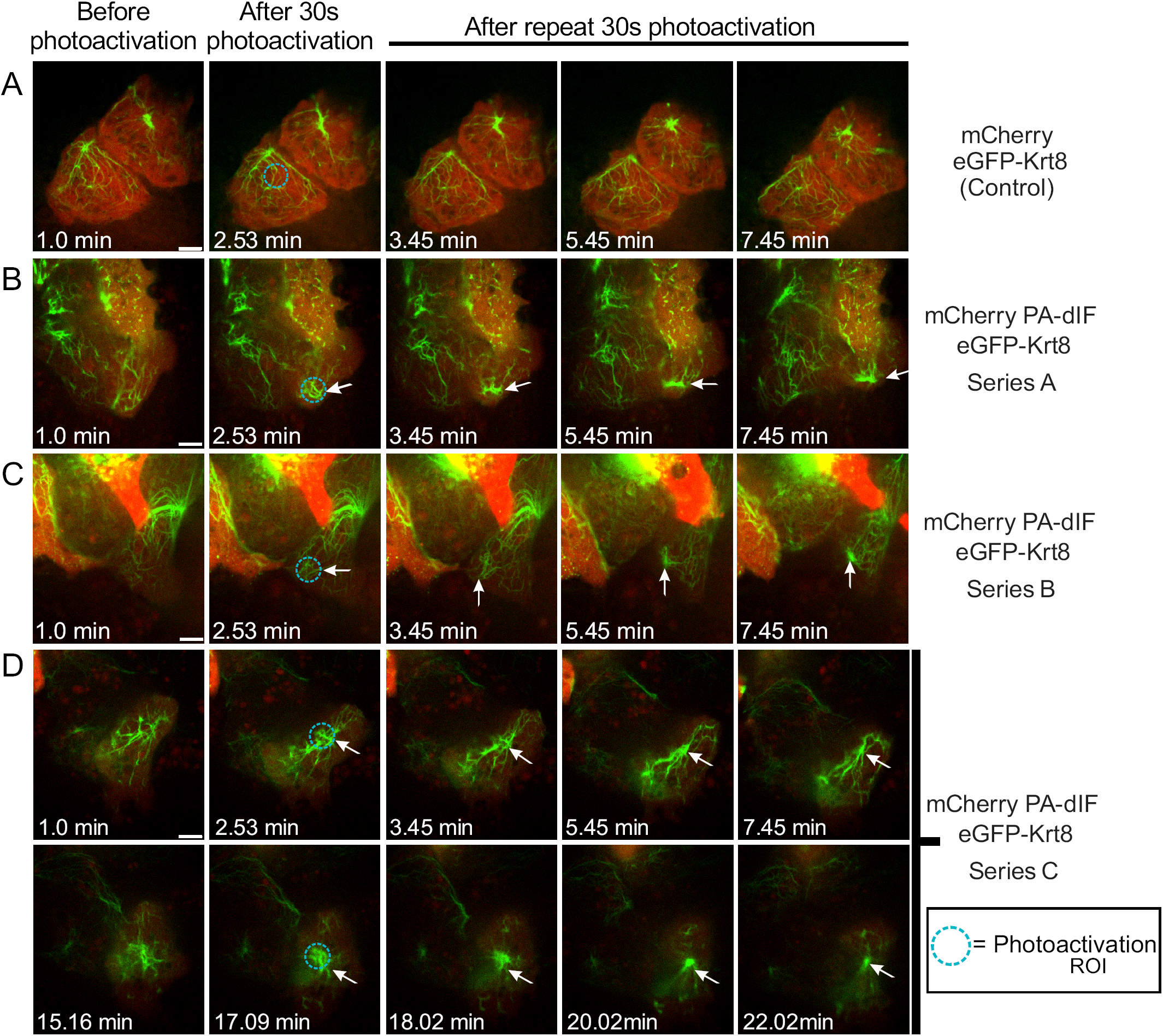
Rapid subcellular disruption of keratin intermediate filaments by photoactivation of a genetically-encoded mimetic peptide. *Xenopus* embryos were co-injected with eGFP-Krt8 and either mCherry or mCherry PA-dIF in the animal cap, targeting the presumptive ectoderm. Cells of the intact ectodermal animal cap explants expressing eGFP-Krt8 (green) and either the control or PA-dIF (red) were illuminated at different subcellular positions with 458 nm light over a 10 µm diameter region (teal circle) for 30s intervals with image acquisition between photoirradiation events. Selected confocal images of time-lapse movies of the response of ectodermal progenitor cells to localized photoirradiation are shown. (**A)** mCherry controls show no keratin disruption nor aggregation either locally or throughout the cell. (**B** and **C)** Two independent examples of light sensitive PA-dIF expressing cells exhibit local disruption and aggregation formation of keratin filaments. (**D)** An extended photoactivation regime induces complete collapse of the keratin network. Arrows indicate region of collapsing keratin network. Circles indicate region where the 458nm laser light was applied. Bars, 10 μm.

### Keratin filaments are necessary for normal *Xenopus* embryo development

Having demonstrated the ability of PA-dIF to disrupt the intermediate filament network in individual cells, we next sought to examine the consequences of intermediate filament disruption on early development. *Xenopus* embryos were injected with mRNA encoding eGFP-Krt19 and mCherry alone, CA-dIF, or PA-dIF and grown to tailbud stages in the dark or in the presence of blue (458 nm) light. Uninjected and mCherry-injected embryos developed normally (Fig. 5, uninjected). In contrast, CA-dIF injected embryos exhibited perturbations in the ectoderm that became apparent during gastrulation (Fig. 5A), a time in which the embryo undergoes major morphogenetic changes as well as increased zygotic protein expression. Extruding yolk plugs could be seen in CA-dIF embryos as they progressed into neurulation (Fig. 5B). By early tailbud and continuing through late tailbud stages, CA-dIF embryos had an open dorsal region, likely originating from the failed blastopore closure (Fig. 5C,D). In tadpoles, this open dorsal aspect persisted in CA-dIF injected embryos (Fig. 5E). Additionally, CA-dIF tadpoles often had underdeveloped head structures, poor midline separation, a bifurcated shortened tail, and a ventral edema (Fig. 5E). These abnormalities were characteristic of CA-dIF/eGFP-Krt19 expressing embryos, but not wildtype or mCherry/eGFP-Krt19 expressing embryos (Fig. 5F).

**Figure 5:**
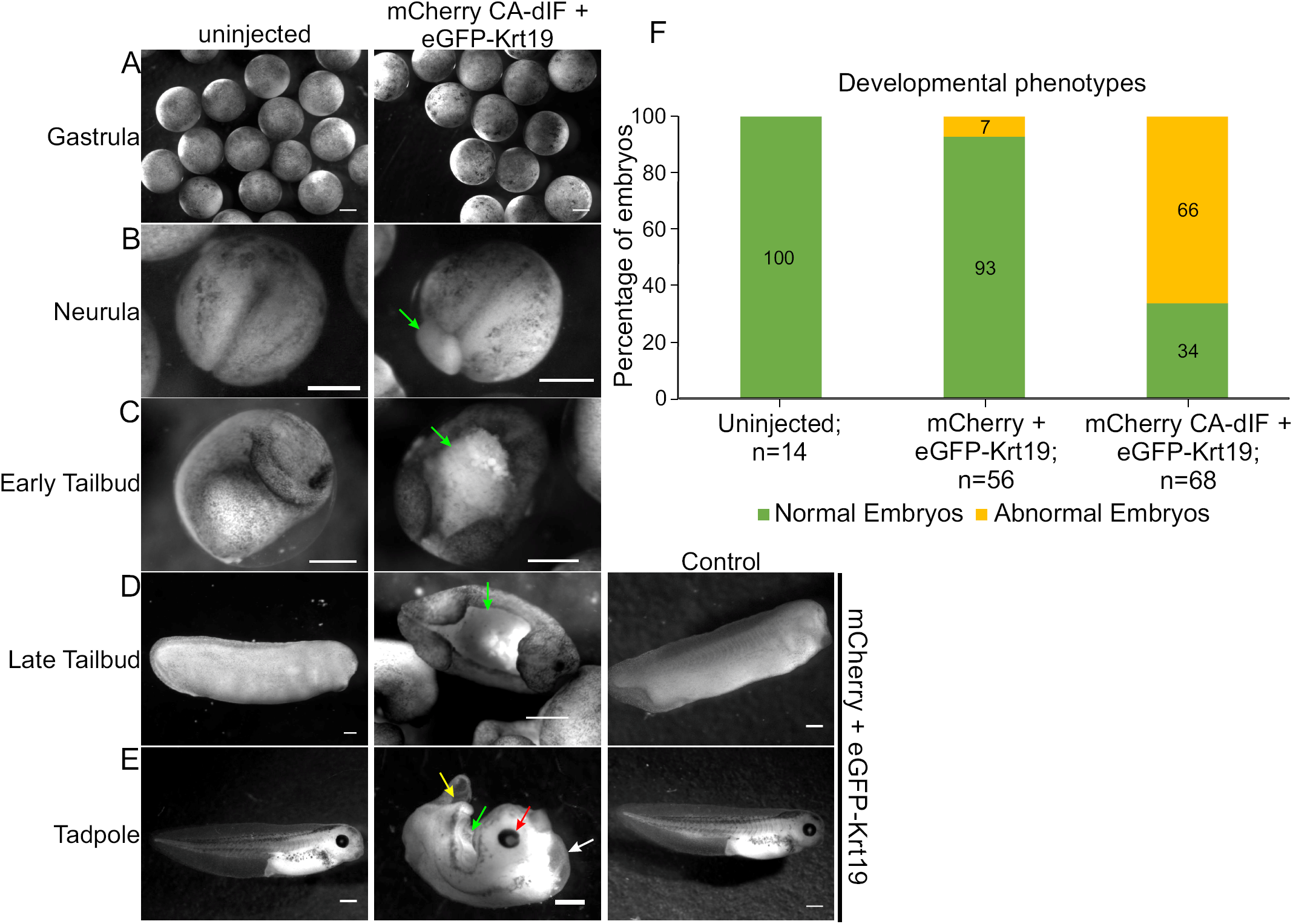
Keratin filaments are necessary for normal *Xenopus* embryo development. Uninjected control embryos or embryos injected with control constructs (mCherry+ eGFP-Krt19) demonstrate normal morphological development from (**A)** gastrulation, (**B)** neurula, (**C)** early tailbud, (**D)** late tailbud, and (**E)** tadpole stages. Expression of mCherry CA-dIF in the ectodermal cells comprising the animal cap induces various developmental defects. Arrows point to defects such as regressed eye development (red arrow), defective yolk plug closure (green arrow), bifurcated tail (yellow arrow), and ventral edema (white arrow). Bars, 500 μm. (**F)** Quantitative representations of normal and abnormal phenotypes.

*Xenopus* embryos injected with 100-175 pg mRNA encoding PA-dIF and maintained in the dark appeared largely like their wildtype and mCherry injected counterparts (Fig. 6A-C). Developmental defects paralleling the CA-dIF phenotypes were induced when PA-dIF embryos were grown under blue light conditions (Fig. 6B,C,F). Injection of greater amounts of PA-dIF mRNA (250-500 pg) resulted in developmental abnormalities irrespective of blue light exposure that were only mildly enhanced by blue light exposure (Fig. 6D-F). Nonetheless, the developmental perturbations due to photoactivation of PA-dIF and its expression at higher levels were consistent with those observed by CA-dIF expression. Altogether, these data show an important role for intermediate filaments during development of the ectoderm in the early embryo.

**Figure 6:**
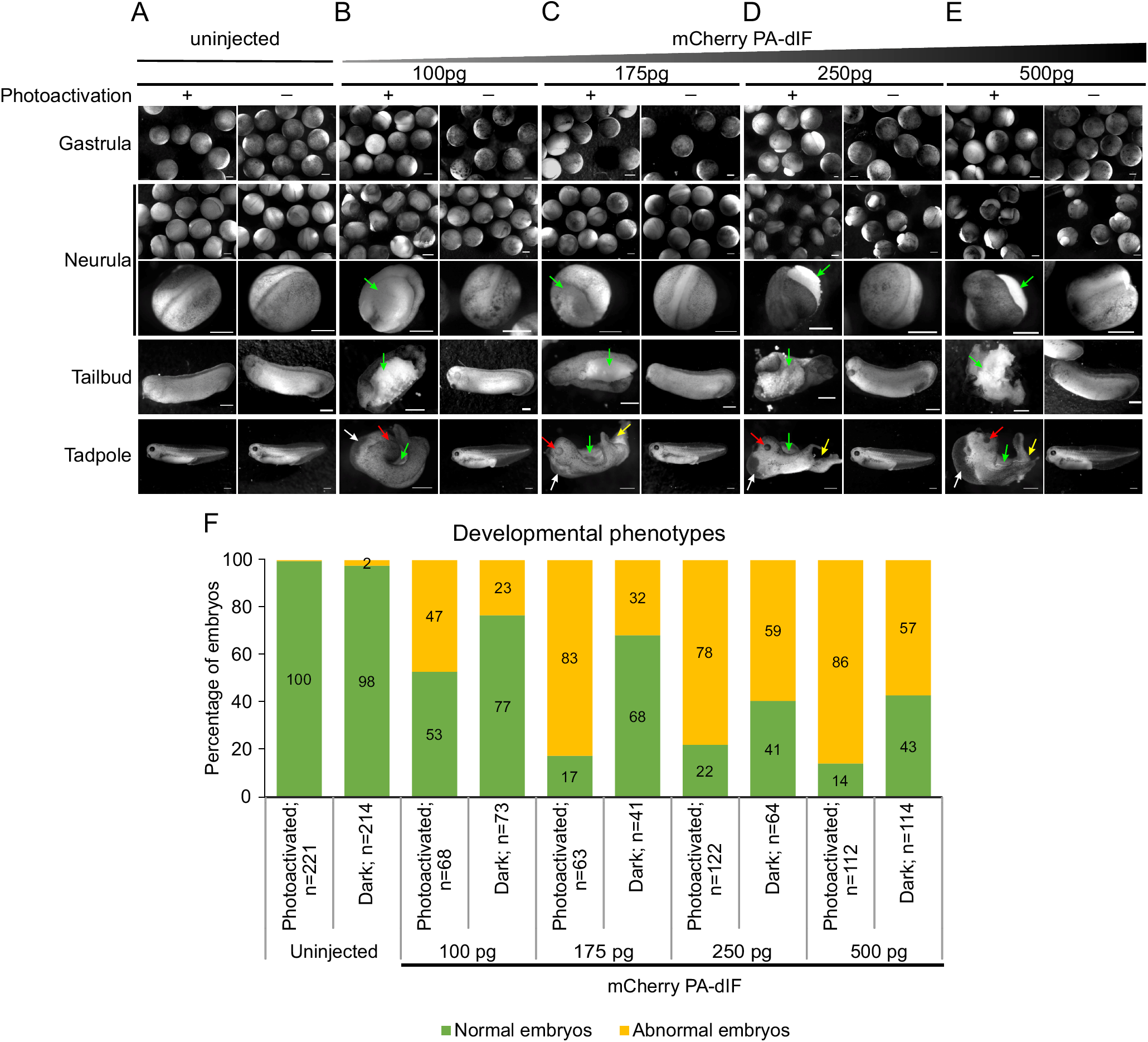
Light induced activation of PA-dIF elicits abnormal *Xenopus* development. Fertilized *Xenopus* eggs uninjected or injected with PA-dIF at one cell stage were either protected from light or exposed to blue light. Phenotypes were assessed from gastrulation to tadpole stage. (**A)** Uninjected control embryos display a normal phenotype regardless of exposure to blue light. (**B)** Embryos were injected with 100 pg PA-dIF, (**C)** 175 pg PA-dIF, (**D)** 250 pg PA-dIF, (**E)** 500 pg PA-dIF. Embryos were grown under blue light conditions as indicated (+ or -). Developmental defects of open yolk plug/open dorsal side (green arrows), under-developed eye or no eye (red arrows), bifurcated tail (yellow arrows), ventral edema (white arrows) are indicated. Bars, 500 μm. (**F)** Quantitative representations of normal and abnormal phenotypes.

## Discussion

We describe herein the generation of PA-dIF, a novel genetically encoded, photoactivatable peptide that can be used to rapidly disrupt intermediate filaments with subcellular spatial resolution. Notably the disruption of the intermediate filaments described is induced without introduction of mutations to intermediate filaments or manipulation of intermediate filament expression levels. We accomplished the disruption by introducing expression of a short peptide that can be made available for disruption through photomanipulation and titered as necessary. Through this approach, these constructs can be used to target intermediate filaments at a subcellular scale or more broadly disrupt in cells and tissues where expressed and photoactivated. In addition, we have used the classic developmental biology system of *Xenopus* and the emergence of developmental phenotypes to test this novel photoactivatable biomolecular tool.

Deciphering the role of keratins in early embryonic development has thus far been difficult, due to functional redundancy and complexity within the family, generation of exogastrulation or embryonic lethal phenotypes in intermediate filament knockout and knockdown animal models, and a lack of specific polymerization inhibitors characterized for intermediate filaments. *In vitro*, 2B2 peptide of vimentin inhibits intermediate filament assembly (Strelkov *et al.*, 2002). This disassembly resulted in formation of unit length filament-like structures. Similarly, synthetic 2B2 peptides based on keratins as a template also interfered with both intermediate filament assembly and disrupted pre-assembled filaments (Hatzfeld and Weber, 1992). *In vivo*, microinjection of 2B2 peptide in fibroblasts causes disruption of vimentin network into short filaments and ULF-like structures (Helfand *et al.*, 2011). In our study using a sequence similar in length to Hatzfeld and Weber (Hatzfeld and Weber, 1992), we observe primarily disruption, collapse and aggregation of the filaments at the site of PA-dIF activation; however, the details of this perturbation and nature of the effects on polymerization remain to be determined. As expected, CA-dIF yields a more widespread disruption and the intermediate filaments appear sparser, less well-defined and often punctate.

Questions remain as to how exactly PA-dIF is incorporating and disrupting the existing filamentous network. We observed that disruption of the intermediate filament network remains sharply within the region of photoactivation. It is apparent that this tool is acutely activated by exposure to blue light resulting in disruption of intermediate filaments. Expression of PA-dIF needs to be tightly regulated and we found that titration of PA-dIF to appropriate expression levels was essential to maintaining dynamic light responsiveness. High expression levels of PA-dIF was sufficient to cause disruption even in dark conditions, similar to CA-dIF. In view of this, improving caging and/or expressing PA-dIF under a titratable expression system will be necessary for greater utility across other cell systems. We have yet to explore whether mutations in LOV-Jα may enhance caging (Zimmerman *et al.*, 2016), but the current photoactivatable construct is sufficient to prevent dIF disruption of the keratin network in the dark and under moderate expression. We also found that CA-dIF can be used to disrupt keratin intermediate filaments simply by expression in cells of multiple species and types. Future work will be aimed at determining the cell biological consequences of intermediate filament perturbation using these tools.

Given the highly conserved sequence homology, we expect that dIF constructs may also disrupt multiple intermediate filament networks. In our LC-MS/MS analyses, vimentin and nuclear lamins were associated with CA-dIF; however, they were found at a much lesser extent and did not meet our stringent proteomics criteria. We did observe notably lobular nuclei, particularly in HEK293T cells transfected with CA-dIF. This nuclear morphology may have resulted from either direct perturbation of nuclear lamins or, more likely, indirect effects on the nucleus that resulted as a consequence of keratin disruption in the cytoplasm. While we cannot determine that the effects of dIF are due exclusively to disruption of the keratin network, this possible affinity of dIF to multiple intermediate filament proteins suggests that the tool may broadly be used to disrupt intermediate filament networks. In this respect, dIF constructs are among a very limited set of tools that can be used for the interrogation of intermediate filaments (Zwerger *et al.*, 2015; Ridge *et al.*, 2016). Nonetheless, Krt18 appeared to be the primary target for our dIF constructs in the early *Xenopus* embryo. Interestingly, we did not find Krt19 particularly enriched in CA-dIF expressing samples even though recent work from our laboratory indicates an important role for 14-3-3 binding to Krt19 for intermediate filament recruitment to cell-cell adhesions (Mariani *et al.*, 2018). Conversely, in that study, we did not find Krt18 associated with 14-3-3. We do not yet know why Krt19 was not associated with our dIF constructs but tentatively speculate that the association of 14-3-3 with Krt19 and/or subcellular compartmentalization may diminish dIF-Krt19 interaction.

Keratin filaments have important roles in normal morphogenesis during *Xenopus* embryonic development as revealed by these new biomolecular tools: PA-dIF and CA-dIF. In prior work, we have inhibited overall keratin expression by injection of embryos with morpholinos, which block translation of zygotic protein. Krt8 morpholino injected embryos exhibited an exogastrulation phenotype characterized by extrusion of the vegetal endoderm out the blastopore (Weber *et al.*, 2012). The phenotypes elicited by morpholino knockdown were similar to those observed in embryos injected with antibodies against keratins (Klymkowsky *et al.*, 1992) or oligonucleotide-induced knockdown of keratin expression (Heasman *et al.*, 1992; Torpey *et al.*, 1992). Here, we describe a third method of disrupting intermediate filaments. Unique to the usage of dIF constructs versus prior knockdown methods is that the present tool distinguishes the role of intermediate filaments as a cytoskeletal network from intermediate filament proteins more generally. In addition, expression of the dIF constructs is more selective for particular tissues because expression is limited to the region proximal to injection (Colman and Drummond, 1986) whereas morpholinos readily diffuse throughout the *Xenopus* embryo (Nutt *et al.*, 2001). In our studies here, we limited expression to presumptive ectodermal tissues through targeted injection of the animal cap. Morphogenetic processes associated with the ectodermal tissue were indeed the main manifestation of the induced defects. Although involution and blastopore constriction have been claimed to be directed primarily by morphogenetic movements in the vegetal and marginal zones (Keller *et al.*, 2003), we induced incomplete yolk plug closure through perturbation of keratins in the ectoderm. Previously it has been suggested that ectodermal movements are an initiating and essential component of subsequent gastrulation movements (Beloussov *et al.*, 2006). Indeed, many perturbations that disrupt ectodermal movements (i.e. epiboly) and extracellular matrix assembly on the blastocoel roof also frequently result in exogastrulation (Rozario *et al.*, 2009; Eagleson *et al.*, 2015). Our data provided here are further experimental evidence that ectodermal movements on their own are an important contributing factor to the overall movements leading to complete blastopore constriction.

Future experiments will determine the contribution of intermediate filaments to morphogenetic movements in other tissues. Furthermore, we now have the capability to examine the role of intermediate filaments in stages beyond gastrulation by simply waiting to expose embryos to blue light until developmental stages of choice. With the flexibility of both spatial and temporal disruption using PA-dIF, the utility of PA-dIF for examining functional developmental roles of intermediate filaments is limitless.

## Supporting information

## Acknowledgements

We would like to thank the members of the Weber laboratory for helpful discussions and advice. Special thanks to Abid Haque of the Weber lab who constructed the blue light box used for the photoactivation of whole embryos. This research was supported by a Rutgers University-Newark Dissertation Fellowship to R.S-S. and grant HD084254 from the Eunice Kennedy Shriver National Institute of Child Health and Human Development to G.F.W. The mass spectrometry data were obtained from an Orbitrap mass spectrometer funded in part by an NIH grant NS046593, for the support of the Rutgers Mass Spectrometry Center for Integrative Neuroscience Research.

## Supplemental figure legends

**Figure S1:**
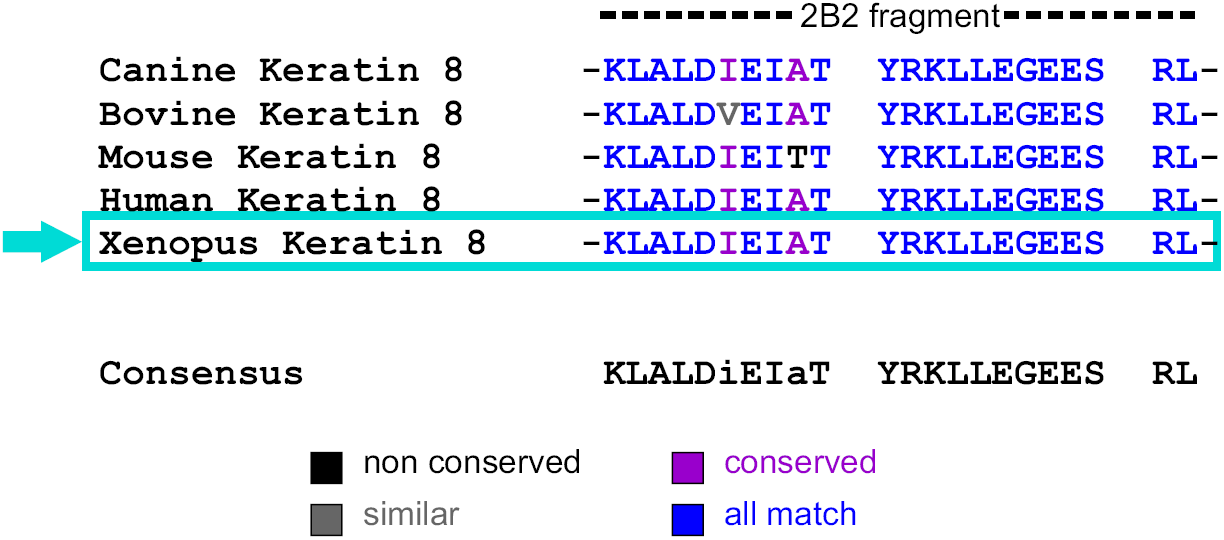
Sequence alignment of 2B2 segments of keratin 8 IF proteins. Protein sequence alignment reveals that the 2B2 peptide is highly conserved across different species including *Xenopus*, Canine, Bovine, Mouse and Human.

**Figure S2:**
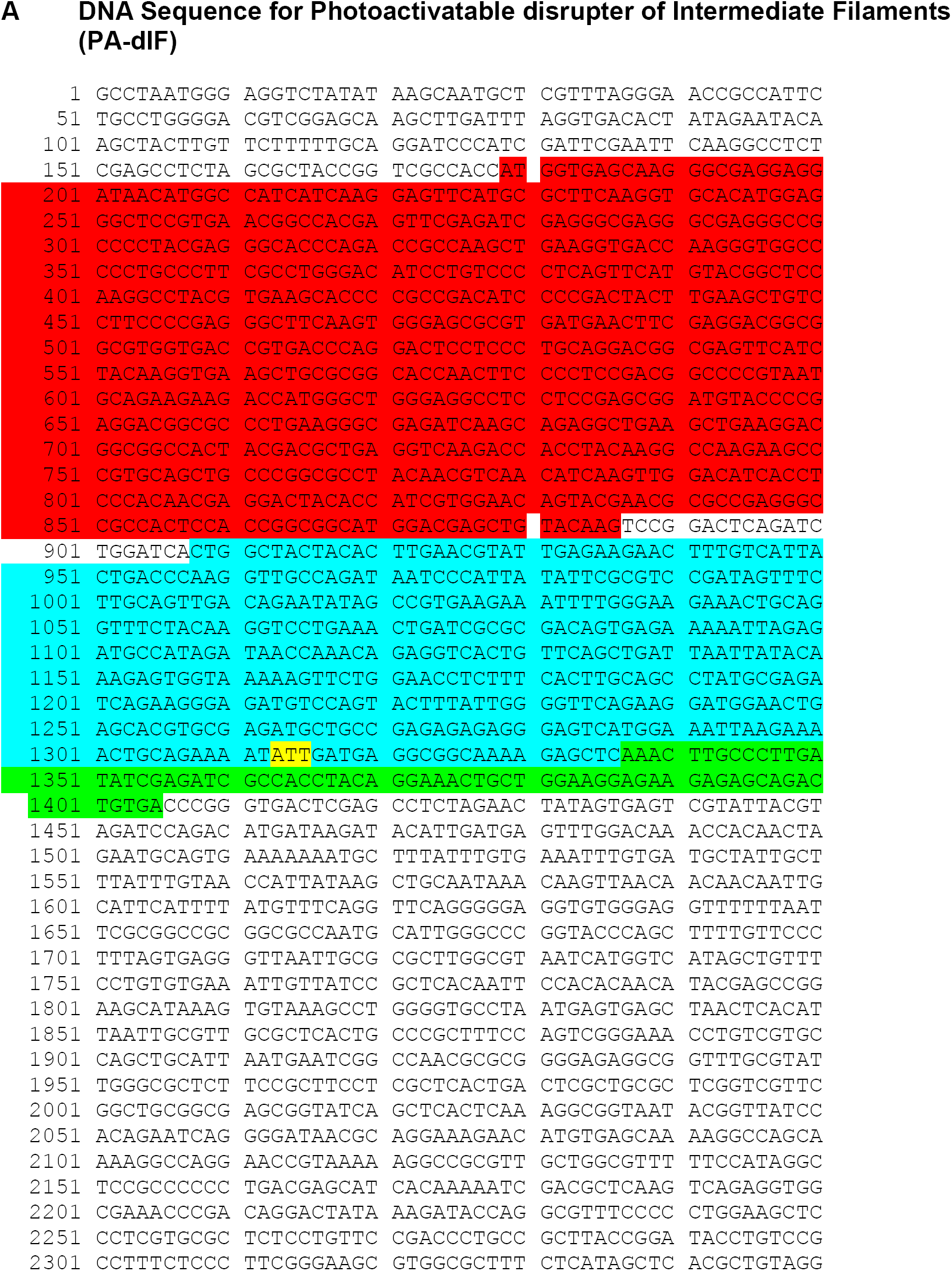
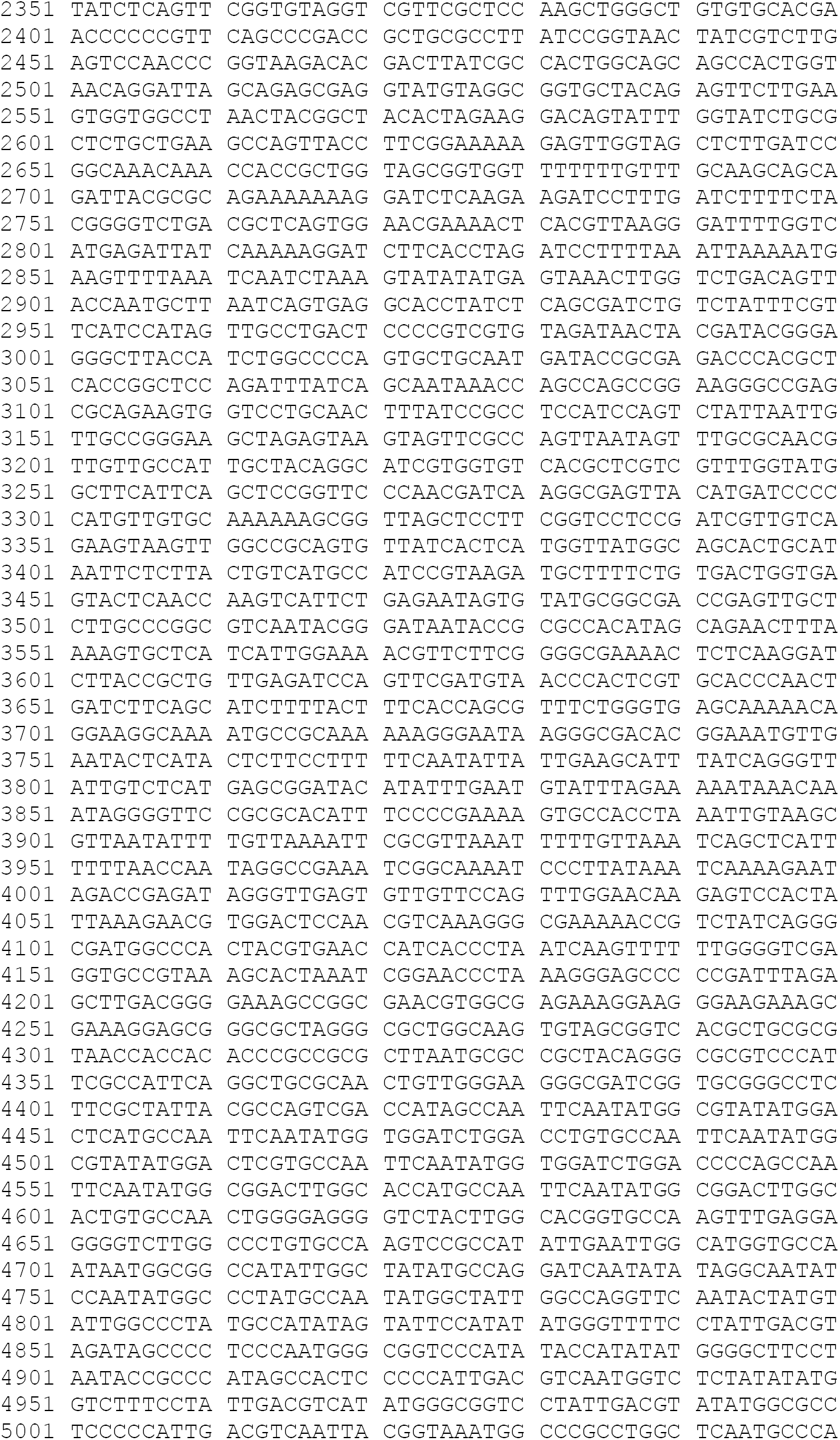
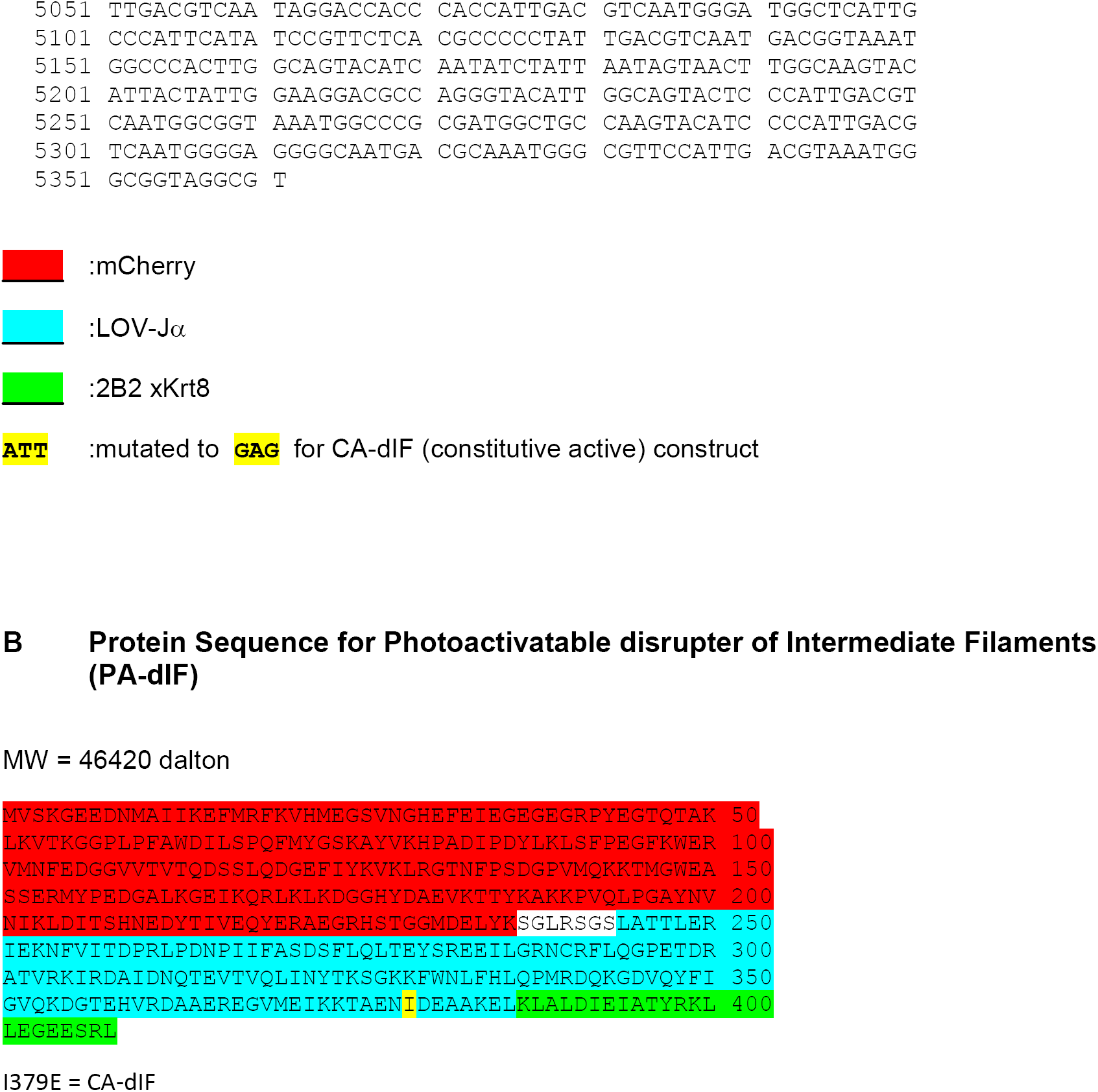
DNA and protein sequences for Photoactivatable disrupter of Intermediate Filaments. DNA sequence for PA-dIF (**A)** and protein sequence for PA-dIF (**B)**. The domains mCherry, LOV-Jα and 2B2 from Krt8 are color coded in red, blue and green respectively.

**Figure S3:**
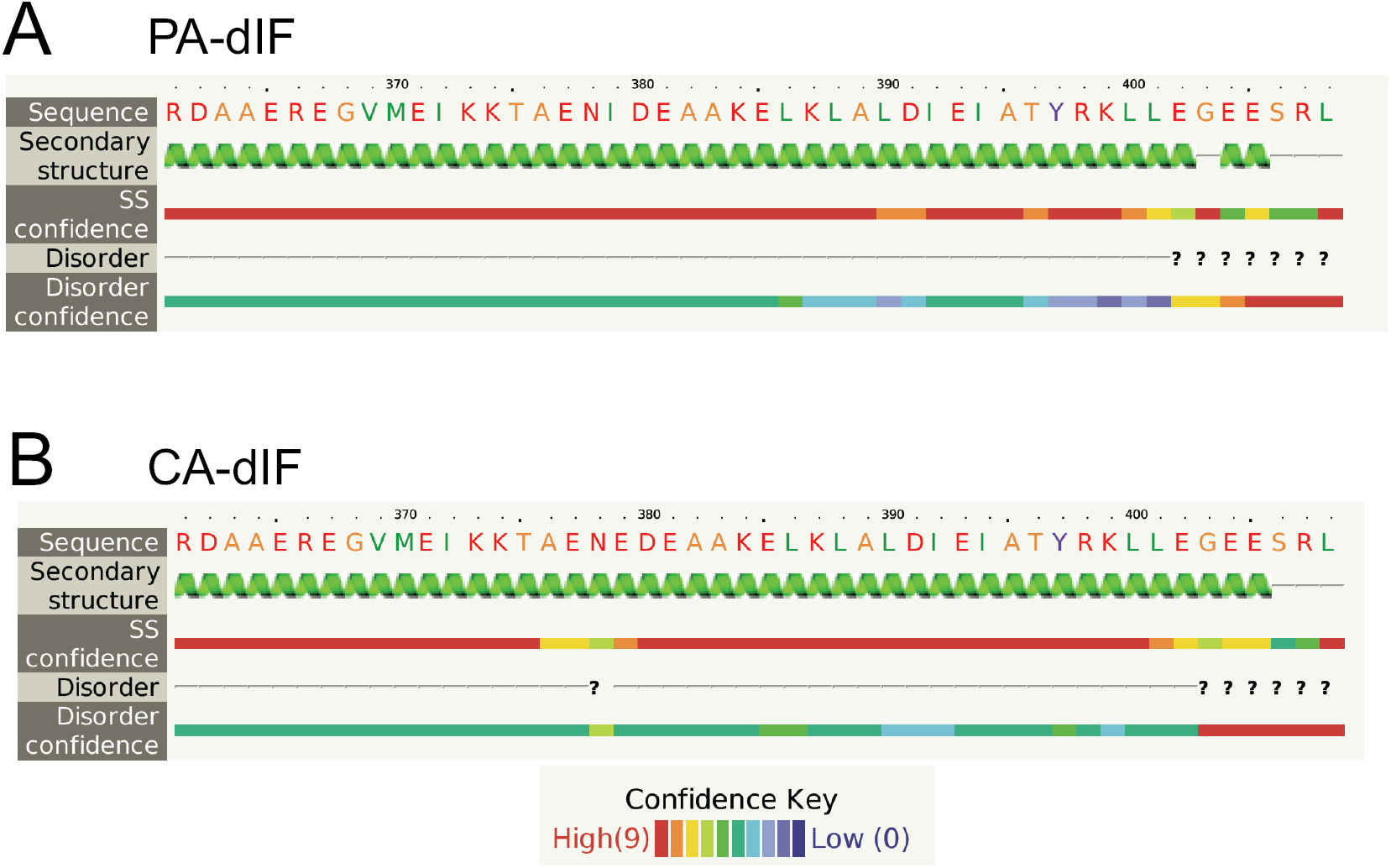
Secondary structure shows continuous α-helical domain. Fusion of Jα to 2B2 retains the α-helical structure in both PA-dIF (**A)** and CA-dIF (**B)**.

**Figure S4:**
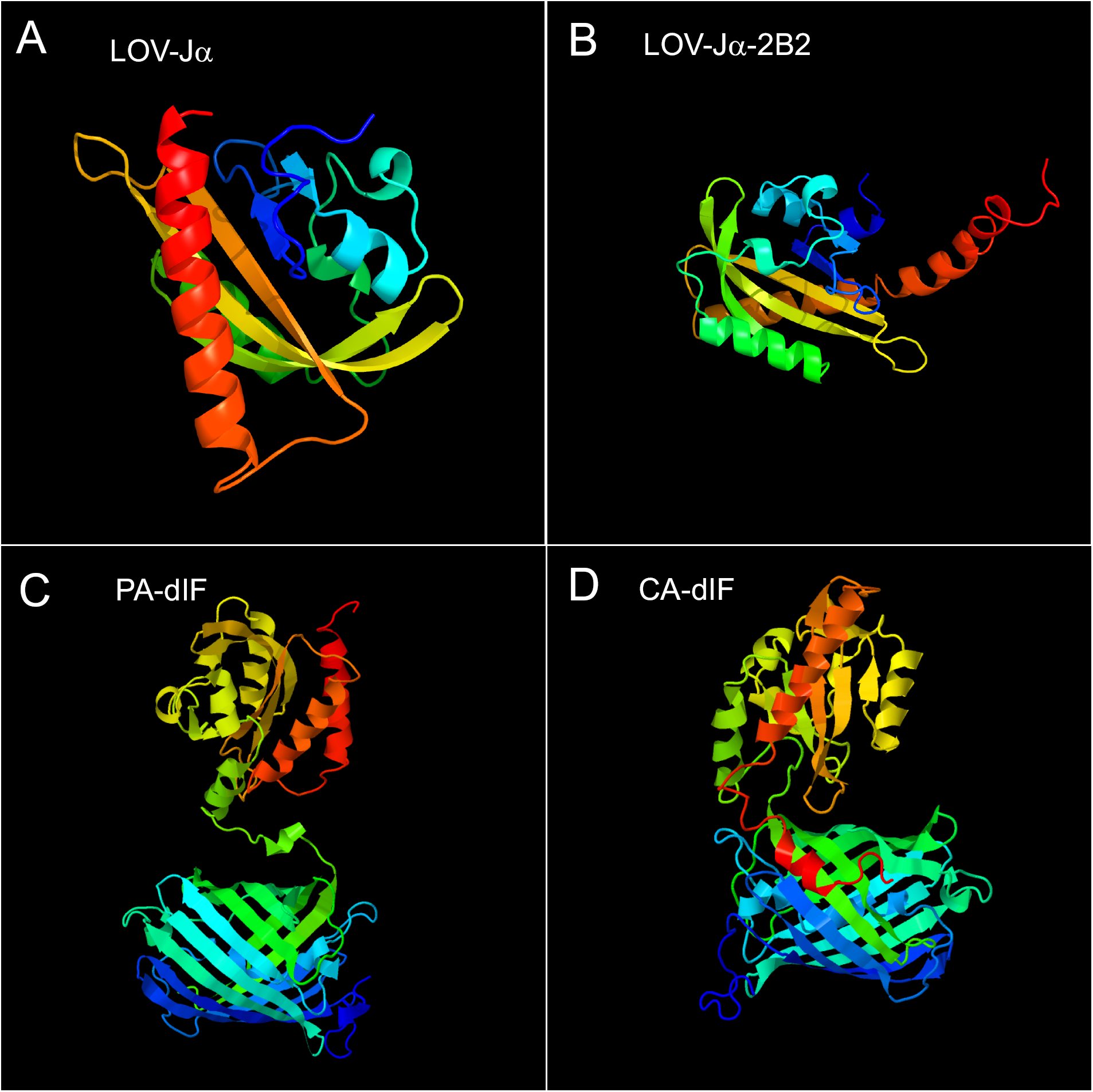
Structural modelling shows a nearly contiguous α-helical domain. In both LOV-Jα (**A)** and LOV-Jα-2B2 (**B)** the 3D structural analysis shows a continuous helical structure of Jα and also of Jα-2B2. However, in PA-dIF (mCherry LOV-Jα-2B2) (**C)**, and CA-dIF (mCherry LOV-Jα(I379E)-2B2) (**D)** the addition of mCherry generates a loop and turn in Jα-2B2. The presence of mutation I379E in (D) induces a steric hindrance of the LOV-Jα interaction not allowing the protein to fold properly. The protein is color coded from N-terminal (blue) to C-terminal (red).

## Supplemental movies

Movie S1: Unperturbed keratin filaments in control ectodermal cells. This movie corresponds to Figure 4A.

Movie S2: Localized photoactivation of PA-dIF induces disruption of keratin filaments-Series A. This movie corresponds to Figure 4B.

Movie S3: Localized photoactivation of PA-dIF induces disruption of keratin filaments-Series B. This movie corresponds to Figure 4C.

Movie S4: Repeated photoactivation of PA-dIF induces complete collapse of the keratin network-Series C. This movie corresponds to Figure 4D.

